# *C. albicans* ergosterol modulates the antifungal response of human neutrophils by masking β-glucan

**DOI:** 10.64898/2026.05.18.721578

**Authors:** Huan Jiang, Angela Nobbs, Ian Leaves, Neil A.R. Gow, Stephanie Diezmann, Borko Amulic

## Abstract

**Introduction:** Ergosterol-targeting azoles are widely used in the treatment of *Candida albicans* infection. In addition to direct antifungal activity, azoles are known to enhance neutrophil-mediated killing of *C. albicans*, but the underlying mechanisms remain unclear, particularly whether ergosterol depletion directly modulates host immune responses.

**Gap Statement:** It remains unknown whether reduced ergosterol levels alone, independent of broader disruption to sterol biosynthesis and fungal morphogenesis, influence neutrophil antifungal activity.

**Aim:** This study aimed to determine how genetic disruption of late-stage ergosterol biosynthesis affects neutrophil-mediated responses to *C. albicans*.

**Methodology:** Doxycycline-repressible GRACE mutants targeting late-stage ergosterol biosynthesis genes (*ERG4, ERG5, ERG3* and *ERG28*) were co-incubated with primary human neutrophils. Fungal survival, oxidative burst, phagocytosis, neutrophil extracellular trap (NET) formation and cell wall composition were assessed.

**Results:** All ergosterol-deficient strains induced elevated neutrophil reactive oxygen species (ROS) production; however, only *ERG4* depletion was associated with enhanced fungal clearance. This phenotype correlated with increased phagocytosis and reduced NET formation. Cell wall analysis revealed no changes in total chitin or mannan content but demonstrated significantly increased surface exposure of β-1,3-glucan in *ERG4*-depleted cells.

**Conclusion:** These findings indicate that disruption of late-stage ergosterol biosynthesis, particularly via *ERG4*, enhances neutrophil antifungal responses and is associated with increased β-glucan exposure. This study highlights a potential role for ergosterol in immune evasion and suggests that targeting terminal steps of the pathway may improve host-mediated clearance of *C. albicans*.

## Introduction

*Candida albicans* is a common component of the human mucosal and skin microbiota, but can also become an opportunistic pathogen^1^. Globally, *C. albicans* is the species most commonly causing invasive candidiasis^2^, and has been categorised as a critical pathogen of the WHO fungal priority pathogen list^3,4^. Approximately 1.5 million people suffer from invasive candidiasis annually, with a 63.6% mortality rate^5^. In addition, at least 150 million cases of recurrent mucosal *Candida* infections are recorded annually^6,7^. The increasing number of immunocompromised patients, including those with prolonged neutropenia or in intensive care, has contributed to a rise in *Candida* infections and antifungal use ^3,8^.

Current first-line treatments for invasive candidiasis, including echinocandins and azoles, are generally effective^2^. Within the azole class, fluconazole remains the primary therapeutic choice, due to its oral bioavailability and broad-spectrum efficacy^9^. However, prolonged and widespread use of azoles has led to an increased frequency of resistant *Candida* isolates in clinical settings^10–12^.

Neutrophils are the first type of leukocyte to arrive at sites of infection, serving as critical first responders in the innate immune response against invading pathogens, such as *C. albicans*^13–15^. They eliminate *Candida* through multiple coordinated mechanisms, including phagocytosis followed by intracellular killing, the production of reactive oxygen species (ROS), and the release of neutrophil extracellular traps (NETs), which immobilise and damage fungal cells^15,16^. Collectively, these responses form a critical first line of defence against invasive candidiasis.

Given the central role of innate immune responses in controlling fungal infections, antifungal drugs that enhance host immunity may provide additional benefit. Azoles target the 14-α-sterol demethylase Cyp51 protein, encoded by the *ERG11* gene^17^. This inhibition disrupts ergosterol biosynthesis, leading to membrane dysfunction and accumulation of toxic sterol intermediates^17^. Apart from direct antifungal mechanisms, it has been reported that azoles act as immunomodulators to enhance the antifungal activity of phagocytic cells^18–21^. For instance, fluconazole has been described to exhibit a synergistic candidacidal effect with human monocytes and monocyte-derived macrophages^18,19^.

In the context of neutrophils, a synergistic effect with azoles has been well documented. Early studies utilising a long-term (24 h) assays reported that while neutrophils alone caused a substantial reduction in the initial fungal inoculum (58-99%), the addition of fluconazole synergised with these cells and enhanced fungicidal activity significantly^22^. Consistently, azoles have been found to interact with cytokine-stimulated human neutrophils, to further promote *C. albicans* killing^23^. For example, posaconazole has been observed to boost neutrophil phagocytic activity and ROS production in response to fungal stimulation^24^.

It remains unclear how ergosterol depletion modulates the immune response. Previous studies demonstrated that *C. albicans ERG11*-deficient cells or cells treated with sub-inhibitory concentrations of clotrimazole becomes hypersensitive to hydrogen peroxide (H_2_O_2_)^25^, suggesting increased susceptibility to oxidative stress. However, it remains unknown whether neutrophil responses are directly enhanced by ergosterol depletion. Disruption of the ergosterol pathway, mediated by either fluconazole treatment or *ERG11* deletion, induces cell wall remodelling characterised by chitin accumulation and β-glucan unmasking, potentially enhancing fungal recognition by the host immune system^26,27^.

A key challenge is that targeting *ERG11* (or early-stage ergosterol synthesis) simultaneously depletes ergosterol and causes morphogenesis defects, both of which independently alter immune recognition^28–31^. To decouple these effects and determine the specific impact of ergosterol deficiency on neutrophil antifungal responses, we used conditional transcriptional repression of enzymes in the late stage of the ergosterol biosynthesis pathway (*ERG4*, *ERG5*, *ERG3*, and *ERG28*). Unlike defects in the early stage, we show that repression of these late-stage genes results in ergosterol depletion whilst cells retain the ability to form hyphae. Through this strategy, we found that *ERG4*-repression significantly increased susceptibility to human neutrophil killing, driven by enhanced phagocytosis and ROS production. Our data further demonstrate that depletion of *ERG4* triggers surface β-glucan unmasking, thereby facilitating effective fungal clearance without initiating the release of neutrophil extracellular traps (NETs).

## Results

### *C. albicans ERG4* and *ERG3* conditional mutants show enhanced susceptibility to neutrophil killing

To avoid the toxicity associated with depletion of early steps of the ergosterol synthesis pathway, we targeted non-essential genes in the late stage of the pathway^32^ (Fig. S1). *ERG4*, *ERG5*, *ERG3*, and *ERG28* were repressed using doxycycline (DOX)-regulated GRACE library mutants, which were subsequently co-incubated *in vitro* with human neutrophils. The transcriptional repression of the target genes was achieved using 0.05 μg/ml DOX (Fig. S2)^28^. Subsequently, *C. albicans* cells were co-incubated with human neutrophils isolated from the peripheral blood of healthy donors. Fungal survival was evaluated using two killing assays: colony forming unit (CFU) enumeration and the Alamar Blue metabolic assays, which measures cellular metabolic activity as a proxy for viability. The survival rates of *ERG5* and *ERG28* conditional mutants were indistinguishable from the wild type (WT) (Fig. 1B, D, F and H). Repression of *ERG3* resulted in significantly increased killing by neutrophils as measured by CFUs, however this effect was not recapitulated in the metabolic assay (Fig. 1C, G). Interestingly, increased susceptibility to neutrophil killing was observed in the *ERG4*-depleted strain in both CFU counting and metabolic activity (Fig. 1A, E). These findings suggest that the final step of ergosterol biosynthesis contributes to fungal survival during neutrophil attack. Together, these data indicated that disruption of the ergosterol pathway does not universally trigger enhanced killing; rather, a strong and consistent susceptibility phenotype appears to be specific to the loss of *ERG4* function.

**Fig. 1.**
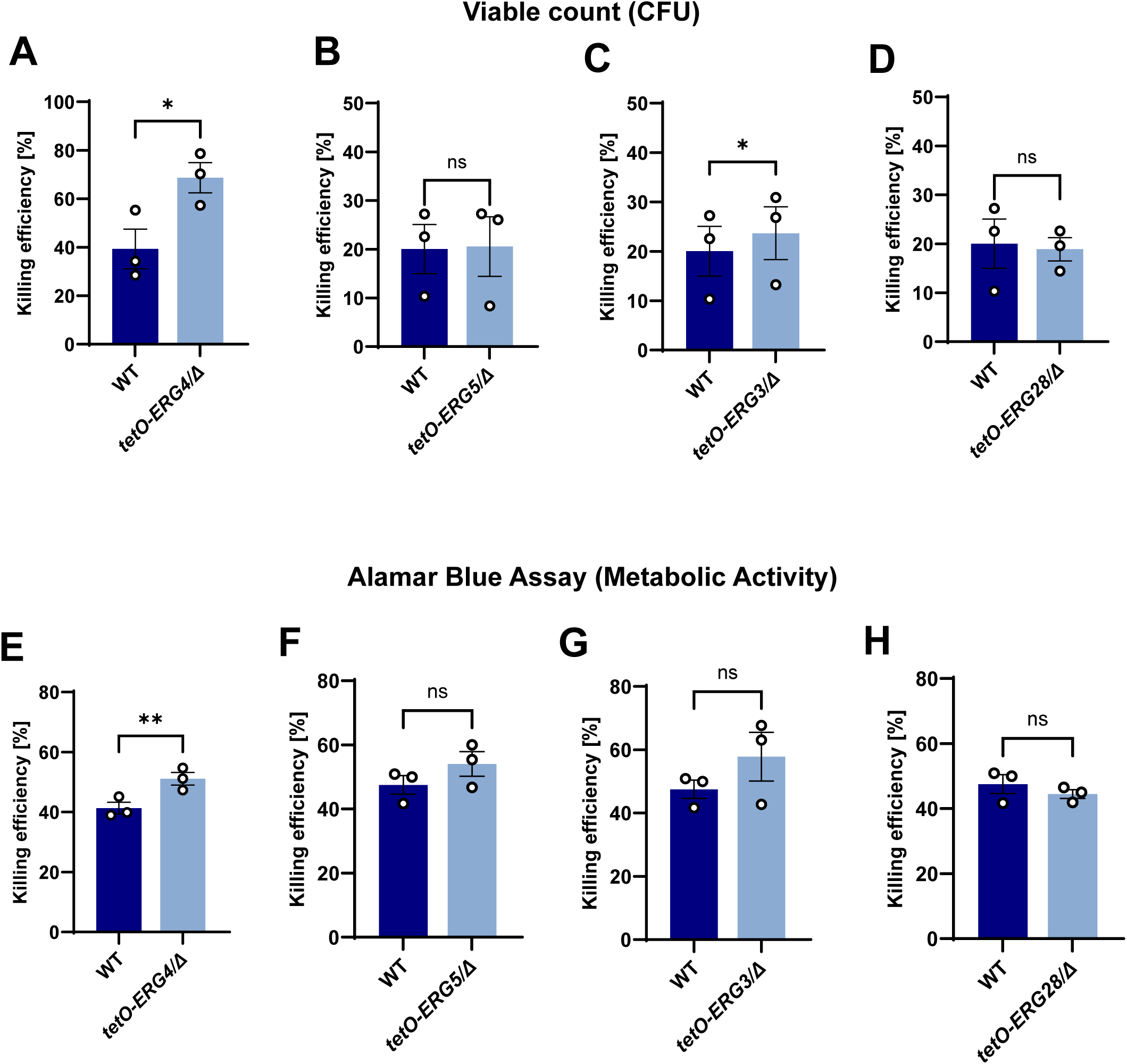
Enhanced neutrophil killing of *C. albicans ERG4* conditional mutant. Human peripheral blood neutrophils were stimulated with *C. albicans* strains at an MOI of 5 in RPMI-1640 with 2% human pooled plasma and 0.05 μg/mL DOX for 2-2.5 h, followed by **(A-D)** plating on YPD agar and incubation for 24 h to obtain viable CFU, or **(E-H)** calculation of metabolic activity measurements from the Alamar Blue assay. Fluorescence signals were normalised to the untreated control, n=3 blood donors, error bars represent mean ± SEM. Significance was calculated using a two-tailed paired Student’s t-test. \**P* < 0.05, \*\**P* < 0.01, ns: *P* > 0.05.

### *C. albicans ERG* conditional mutants show normal growth kinetics and morphogenesis

To characterise the effect of depletion of *ERG4*, *ERG5*, *ERG3* and *ERG28* on fungal growth, we monitored the growth kinetics of the WT and mutant *C. albicans* strains in YPD medium at 30°C for 20 h. Results showed no significant differences in growth (Fig. 2A-D).

**Fig. 2.**
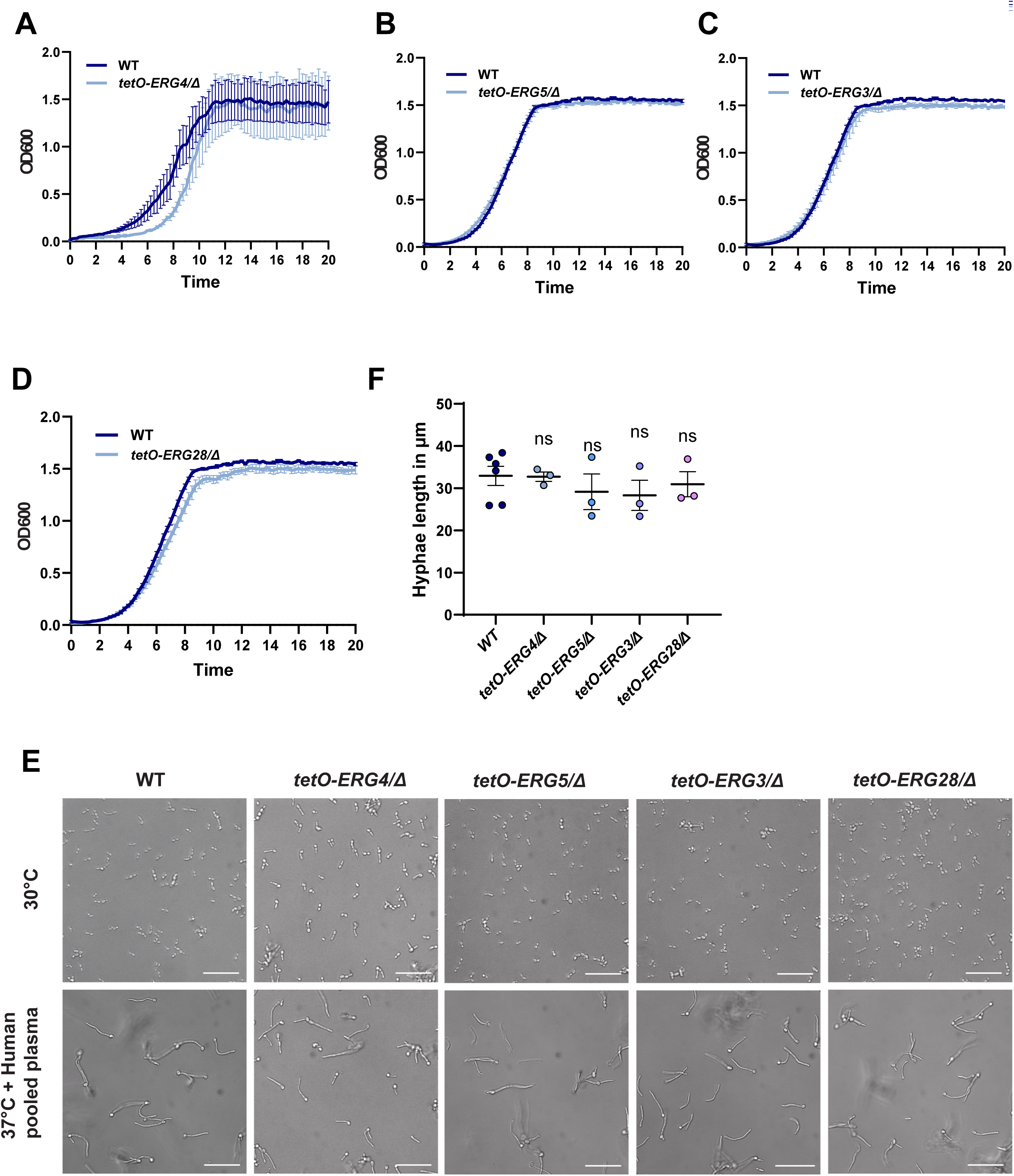
***C. albicans ERG* conditional mutant strains have *in vitro* normal growth kinetics and morphogenesis. (A-D)** Growth kinetics of *C. albicans* strains in YPD at 30°C in the presence of 0.05 μg/mL DOX with starting OD_600_ of 0.02. The OD_600_ of each sample was measured every 15 min over 20 h. **(E)** Hyphal formation of WT and *ERG* conditional mutants induced for 2.5 h in RPMI-1640 with 0.05 μg/mL DOX at 30°C and 37°C (supplemented with 10% human pooled plasma). Representative images of WT, *ERG4-*, *ERG3-*, *ERG5-*, and *ERG28*-depleted *C. albicans* strains at 20X magnification. Scale bar = 50 μm. **(F)** Quantification of average hyphae length (μm) from **(E),** n=3 biological replicates, error bars represent mean ± SEM. Significance was calculated using one-way ANOVA with Dunnett’s multiple comparison post-hoc test. ns: *P* > 0.05.

Neutrophil effector functions are meticulously regulated by distinct *C. albicans* morphotypes in a dose-and surface area-dependent manner^30^. To ensure that the observed killing phenotypes were not driven by morphogenesis defects, we monitored the morphology of each mutant strain in response to hyphal induction with RPMI-1640 medium supplemented with 10% human pooled plasma at 37°C. Quantification of hyphae length after 2.5 h incubation showed no significant effect of transcriptional repression of *ERG4*, *ERG5*, *ERG3*, and *ERG28* on filamentation (Fig. 2E, F). Together, these observations indicate that the susceptibility of *ERG3-and ERG4*-depleted strains to neutrophil killing was not caused by growth or filamentation defects.

### *C. albicans ERG* conditional mutants induce elevated neutrophil ROS production

The respiratory burst is a hallmark antifungal effector mechanism whereby neutrophils produce a rapid and massive amount of ROS with microbicidal properties^15,33^. To determine whether the *ERG* conditional mutant cells were able to modulate ROS production, we utilised a luminol-amplified chemiluminescence kinetic assay. Neutrophils stimulated with WT and mutant strains triggered ROS within 2-5 min, reaching the maximum ROS level by 20 min (Fig. 3A-D). Area under the curve (AUC) analysis demonstrated significantly elevated total ROS production in all *ERG* conditional mutant strains compared to WT, with an average increase of approximately 50% (Fig. 3E-H). In conclusion, genetic deficiency in ergosterol biosynthesis increases the magnitude of the neutrophil oxidative burst.

**Fig. 3.**
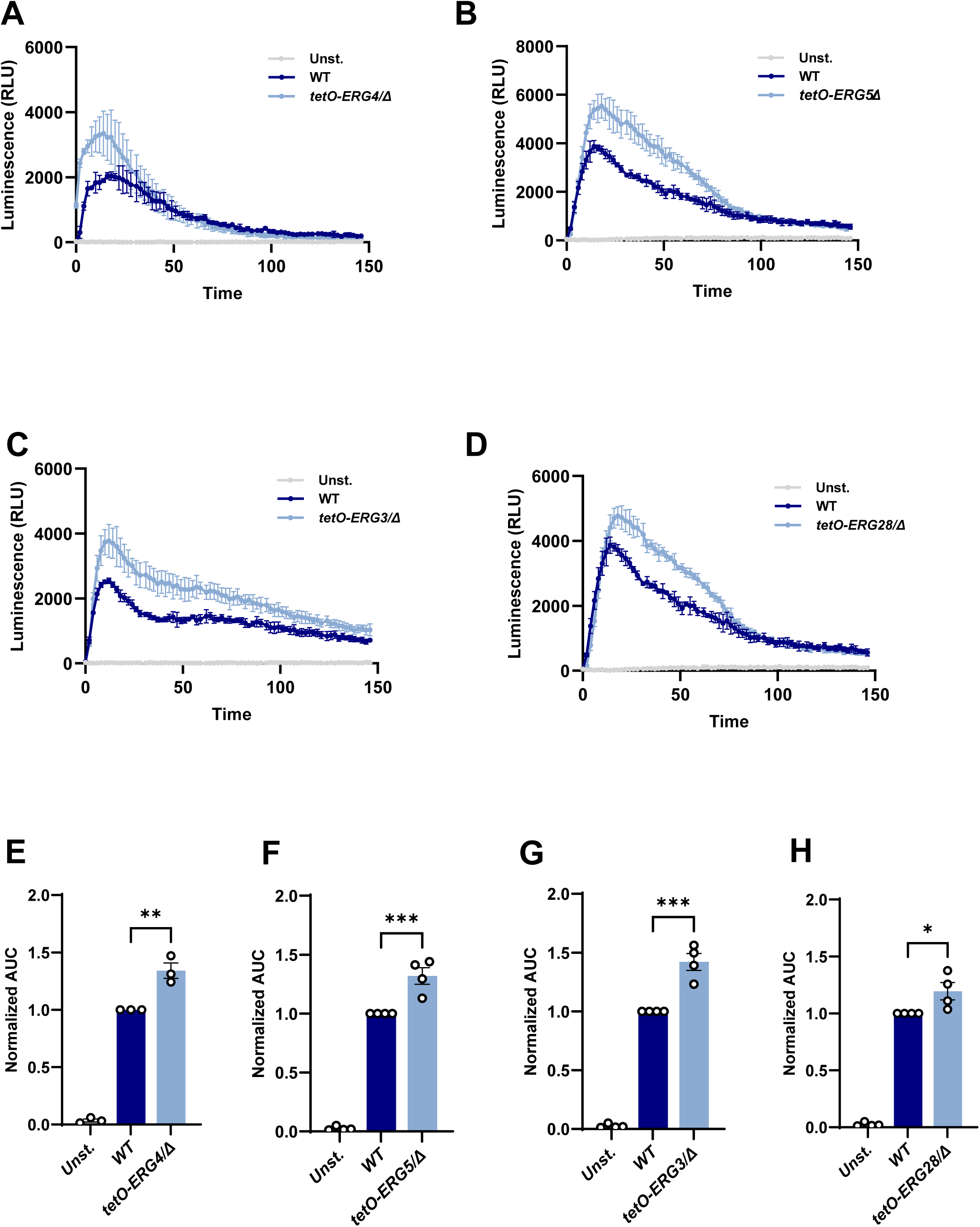
**Elevated neutrophil ROS production in response to *C. albicans ERG* conditional mutants**. **(A-D)** Representative kinetics of the neutrophil oxidative burst, detected with luminol, upon co-incubation with WT, *ERG4-*, *ERG5-*, *ERG3-*, and *ERG28*-depleted *C. albicans* cell (MOI of 5) over 2.5 h at 37°C. As a negative control, neutrophils were left unstimulated. **(E-H)** Quantification of total ROS area under the curve of three independent donors. Values were normalised to the AUC of the WT strain corresponding donors. Error bars represent mean ± SEM. Significance was calculated using one-way ANOVA with Dunnett’s multiple comparison post-hoc test. \**P* < 0.05, \*\**P* < 0.01, ***P < 0.001, ns: *P* > 0.05.

### *ERG4* depletion results in more efficient phagocytotic uptake

To determine whether the enhanced killing and oxidative burst observed in ergosterol synthesis mutants were driven by increased yeast internalisation, we next examined susceptibility to opsonic phagocytosis. Engulfment of yeast cells was assessed by fluorescence microscopy after 30 min co-culture with neutrophils. Internalised fungi were distinguished from external cells via Calcofluor White (CFW) exclusion to calculate the phagocytic index (number of intracellular yeast cells per neutrophil) (Fig. 4A). Depletion of *ERG4* or *ERG5* significantly enhanced phagocytosis, resulting in a nearly 1.5-fold increase compared to WT (Fig. 4B, C). Phagocytosis of the *ERG3-*and *ERG28-*repressed cells demonstrated a non-significant upward trend (Fig. 4D, E), suggesting that the ability to evade neutrophil engulfment is promoted by specific sterol structures in the final steps of ergosterol synthesis catalysed by Erg4 and Erg5.

**Fig. 4.**
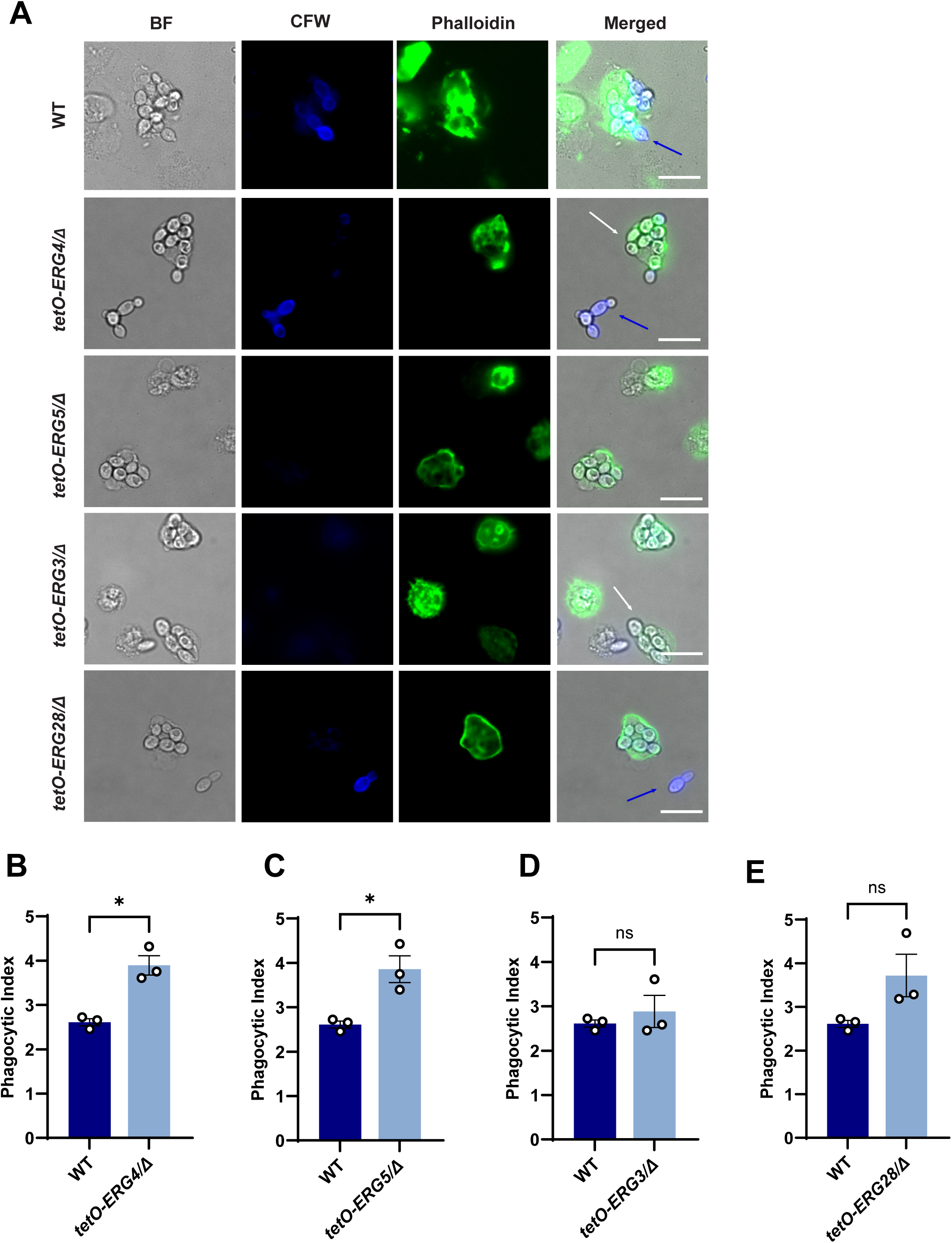
Enhanced neutrophil phagocytosis of strains deficient in late stage ergosterol synthesis. Human peripheral blood neutrophils were stimulated with WT, *ERG4-*, *ERG5-*, *ERG3-*, and *ERG28*-depleted *C. albicans* cells at an MOI of 3 for 30 min. Extracellular *C. albicans* cells were stained with CFW (blue), and neutrophils were labelled with phalloidin (green). **(A)** Representative images of phagocytosed WT and mutant strains. Scale bar = 15 μm. **(B-E)** The phagocytic index was determined as the ratio of the total number of engulfed *C. albicans* cells (unstained) to the total number of counted neutrophils, n=3 blood donors, error bars represent mean ± SEM. Significance was calculated using a two-tailed paired Student’s t-test. \**P* < 0.05, ns: *P* > 0.05.

In contrast, cytokine production remained comparable to WT control (Fig. S3), demonstrating some specificity in ergosterol immune modulation.

### *ERG4* deficiency attenuates NET formation

As an alternative effector killing mechanism, particularly when phagocytosis is insufficient to control large *C. albicans* hyphae, neutrophils release NETs through chromatin externalisation^16,34^. This is a costly response, as NETs are pro-inflammatory and cytotoxic, with a major potential for causing immunopathology^15^. We investigated whether ergosterol depletion affects NET release. Neutrophils were stimulated with WT and *ERG*-depleted *C. albicans* strains for 4 h, after which NETs were visualised by immunofluorescence microscopy with an antibody targeting the specific NET antigen cleaved histone H3 ^35^ (Fig. 5A). Exposure to phorbol 12-myristate 13-acetate (PMA), a standard NETosis inducer, was used as a positive control and elicited extensive extracellular chromatin release compared to unstimulated neutrophils (Fig. 5A). Quantification of the relative NETs area revealed that *ERG4*-depleted cells induced significantly fewer NETs, with nearly 30% reduction compared with the WT strain (Fig. 5B). We also observed a trend for reduced relative NETs area for neutrophils stimulated with the *ERG5* and *ERG3* conditional mutants, but not in the *ERG28*-depleted condition (Fig. 5C, D, E). In summary, these data demonstrated that *ERG4*-mediated sterol modification may promote NET release, whereas depletion of *ERG4* enhances phagocytic clearance over extracellular trap release.

**Fig. 5.**
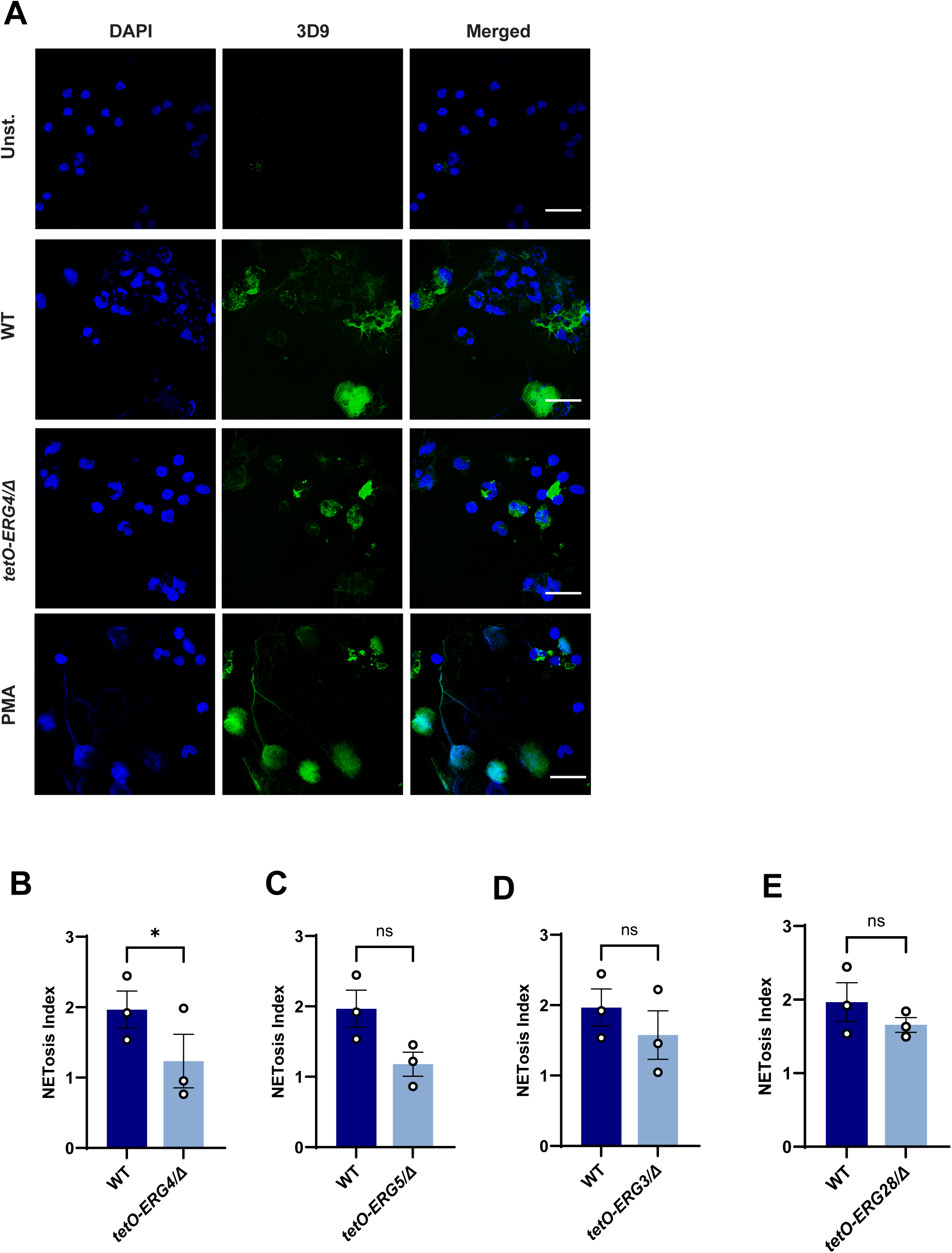
The *ERG4* conditional mutant triggers reduced NET formation. Human peripheral blood neutrophils were stimulated with WT, *ERG4-*, *ERG5-*, *ERG3-*, and *ERG28*-depleted *C. albicans* cells at an MOI of 5 for 4 h. **(A)** Representative confocal images (63X) show DAPI (blue) and NETs visualised by cleaved histone H3 (3D9; green). Scale bar = 20 μm. **(B-E)** Quantification of relative NETs area per 40X field of view. The relative NETs area was quantified as the ratio of 3D9-positive area to DAPI-stained area, averaged across at 6-7 fields of view, (n=3 blood donors), error bars represent mean ± SEM. Significance was calculated using a two-tailed paired Student’s t-test. \**P* < 0.05, ns: *P* > 0.05.

### *ERG4* depletion increases exposure of immunogenic **β**-1,3-glucans

Given the importance of pathogen-associated molecular patterns (PAMPs) in neutrophil activation^33,36^, we assessed whether *ERG4*-depleted mutant displayed major alterations in cell wall composition. The *C. albicans* yeast cell wall consists of an outer layer of mannan and an inner layer enriched in β-1,3-glucans and underlying chitin^37^, which can be recognised by neutrophil pattern recognition receptors (PRRs)^36^. We first assessed whether the global cell wall monosaccharide composition was altered, by performing high-performance ion chromatography (HPIC) analysis, which revealed no significant differences between the WT and *ERG4*-depleted mutant strains (Fig. 6A-C).

**Fig. 6.**
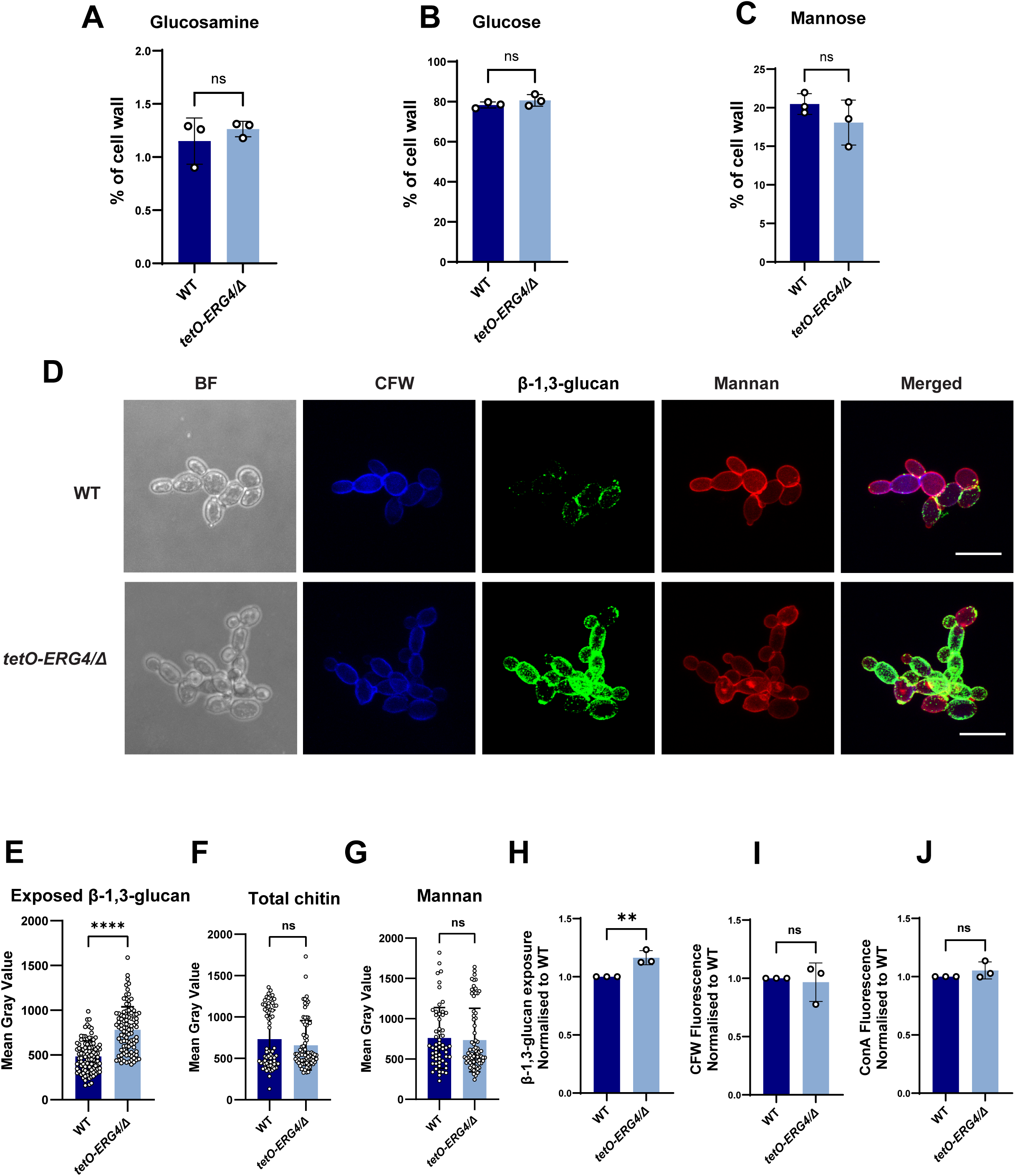
Depletion of the *ERG4* gene increases exposure of β-glucan on the fungal cell wall. HPIC analysis of cell wall **(A)** glucosamine, **(B)** glucose and **(C)** mannose composition in WT and *ERG4*-depleted strain. **(D)** Representative confocal micrographs of mid-log phase cultures of WT and *ERG4*-depleted strains stained with a human Fc-Dectin-1 fusion protein ((β-1,3-glucan), calcofluor white (CFW; chitin), and Alexa Fluor 488 conjugated concanavalin A (ConA; mannan). Scale bar, 15 μm. **(E-G)** Quantification of the relative mean fluorescence intensity in WT and *ERG4*-depleted strains from **(D)**. Each dot represents a field of view (n=3 biological replicates). **(H-J)** Quantification of the mean fluorescence intensity (MFI) levels of exposed β-1,3-glucan, total chitin and mannan by flow cytometry. MFI was normalised to stained WT values of each experimental repeat (n=3). Error bars represent mean ± SEM. Significance was calculated using a two-tailed unpaired Student’s t-test. \*\**P* < 0.01, \*\*\*\**P* < 0.0001 ns: *P* > 0.05.

We hypothesised that the altered neutrophil responses to the *ERG4*-depleted mutants were due to unmasking of immunogenic PAMPs. To assess changes in cell wall epitopes, cells were stained with CFW, concanavalin A (ConA) and human Fc-dectin-1 and to quantify total chitin, mannan and surface exposed β-1,3-glucan, respectively. We used both fluorescence microscopy and flow cytometry to quantify this binding. The abundance of total chitin and mannan remained similar in WT and *ERG4-*depleted yeast cells, as shown with ConA and CFW (Fig. 6F, G, I, J). In WT cells, microscopy showed that Fc-dectin-1 primarily stained the bud scar and bud neck, whereas the mutant exhibited a marked increase in β-1,3-glucan exposure throughout the entire cell wall (Fig. 6D). Quantification of confocal micrographs demonstrated significantly higher levels of β-1,3-glucan on the *ERG4-*depleted mutant, with a 1.5-fold enhanced exposure compared with WT strain (Fig. 6E). Consistent with this, flow cytometry revealed a significant increase in β-1,3-glucan levels in the conditional mutant compared to WT (Fig. 6H). Taken together, these results demonstrated that the depletion of *ERG4* increases surface of immunogenic β-1,3-glucan exposure.

## Discussion

This study shows that disruption of the last stage of ergosterol biosynthesis, through *ERG4* depletion in *C. albicans* exerts direct immunomodulatory effects that boost neutrophil-mediated fungal clearance. Notably, *ERG4* depletion did not impair fungal growth or yeast-to-hyphae morphogenesis in response to human pooled plasma and temperature shift. This is in contrast to the inhibition of Erg11, which disrupts earlier steps in the pathway and leads to toxic sterol accumulation and severe membrane dysfunction^38–41^. Instead, *ERG4* depletion preserved fungal viability whilst altering membrane and cell wall architecture, a phenotype previously characterised in other fungi^32,42,43^.

Transcriptional repression of *ERG4* enhanced oxidative burst and phagocytosis, while suppressing NETs formation. These effects are most likely driven by the unmasking of immunogenic cell wall β-1,3-glucan, causing increased susceptibility to neutrophil fungicidal activity. In contrast, no significant differences were observed in total chitin and mannan levels. However, while chitin and mannan appeared unchanged, we cannot exclude that more subtle alterations in their organisation or surface accessibility contributed to immune recognition. This remains to be investigated.

We observed differences when comparing the *ERG4*-depleted strain with other late stage ergosterol mutants. Although the *ERG5*-, *ERG3*-, and *ERG28*-depleted strains induced similarly elevated ROS levels, they maintained equivalent rates of neutrophil killing. Furthermore, depletion of *ERG3* and *ERG28* did not alter phagocytosis, while the *ERG5*-repressed strain displayed increased uptake without enhancing fungal clearance. These findings indicate that ROS induction alone does not solely drive fungal killing and shows that *ERG4* depletion is specifically associated with increased fungal susceptibility to neutrophil attack. The reason for this discrepancy cannot be elucidated at this point. We hypothesise that the sterol intermediates in *ERG4*-depleted cells, particularly the accumulation of ergosta-5,7,22,24(28)-tetraenol (ET), may disrupt membrane integrity more profoundly than intermediates accumulating in other mutants. Alternatively, *ERG5*-, *ERG3*-, and *ERG28*-deficient strains may initiate compensatory stress response pathways, such as antioxidant defences, allowing them to withstand neutrophil-derived ROS.

*ERG4* encodes a C-24 sterol reductase that converts the precursor ET into the essential component ergosterol^38^. Previous studies in different fungal species supported a critical role for *ERG4* in stress adaptation^42,43^. These findings support the notion that *ERG4* is dispensable for growth but essential for maintaining membrane robustness under stress conditions. In this context, the efficient clearance of the *ERG4* mutant may reflect a dual effect: increased neutrophil activation combined with intrinsic fungal vulnerability to ROS. This proposed mechanism warrants further experimental validation.

Interestingly, our data show increased phagocytosis but reduced NETosis in the *ERG4* conditional strain. NETs preferentially facilitate the killing of microbes that cannot be efficiently engulfed by phagocytosis, such as hyphae^44–46^. In fact, Branzk et al. also showed that successful Dectin-1 mediated phagocytosis actively downregulates NETosis by sequestering neutrophil elastase within the phagosome^46^. Consistent with this model, the rapid and efficient phagocytosis of *ERG4*-depleted cells likely reduces the requirement for extracellular trap release, indicating a functional trade-off between internalisation and NETs deployment. This may have therapeutic advantages, as limiting NETosis could reduce host immunopathology^15^.

Neutrophil detection of *C. albicans* is thought to rely on opsonic and non-opsonic pathways. We included human pooled plasma in all our assays, meaning both pathways are relevant. β-glucan can be directly detected via complement receptor 3 (CR3) and C-type lectin receptor Dectin-1, which drives downstream antifungal responses^36,47^. Activation of Dectin-1 promotes phagocytosis, consistent with the immunomodulatory role of exposed β-glucan described in previous studies^36,48^. Cell wall remodelling following *ERG* depletion appears to be a conserved feature across fungi. For example, in *Cryptococcus neoformans*, increased surface-exposed chitin has been reported as a compensatory structural response to membrane stress^42^. Given that our current study quantified total chitin levels, further assessment of surface-exposed chitin is necessary to determine if a similar compensatory remodelling occurs in *C. albicans*.

In conclusion, this study reveals that ergosterol biosynthesis in *C. albicans* contributes to immune evasion by masking β-glucan from neutrophil recognition. This positions *ERG4* as a novel immune evasion gene. Our findings identify *ERG4* as a key immunomodulatory target that enhances fungal clearance without compromising viability, suggesting that targeting late stage ergosterol synthesis may represent a viable therapeutic strategy. Future studies should explore the *in vivo* potential and evaluate whether targeting Erg4 renders azole treatment curative rather than merely fungistatic.

## Materials and methods

### *C. albicans* growth conditions

The *C. albicans* GRACE library was obtained from the National Research Council of Canada^49^. Yeast cells were cultured for 16 h at 30 °C, 200 RPM in YPD medium (1% yeast extract, 2% peptone from meat). Suppression of target gene expression in GRACE strains was induced by adding the indicated concentrations of DOX to the growth medium. All stocks were preserved at −80 °C in 25% glycerol.

### Primary neutrophil isolation from peripheral blood

Blood samples were collected from consenting healthy donors at the University of Bristol following ethical approval from the NHS Research Ethics Committee (REC 18/EE/0265). The samples were drawn into EDTA tubes for collection. Neutrophil isolation was performed using the negative selection EasySep direct human neutrophil isolation kit (STEMCELL Technologies, Cambridge) as per the manufacturer’s instructions.

### *C. albicans* reverse transcription and q-PCR

Strains were grown in YPD at 30°C in the presence or absence of 0.05 μg/ml DOX for 16 h, diluted to an OD_600_ of 0.1 then grown to mid-log phase for 3 h. Pellets were stored at-80°C pitot to RNA extraction. RNA was isolated using the RNeasy Mini kit. Complementary DNA strand synthesis was conducted using the AffinityScript QPCR cDNA Synthesis Kit (Agilent Technologies). qRT-PCR was performed using FAST SYBR Green Master Mix (Thermo Fisher Scientific) and Applied Biosystem Real-Time PCR Instruments, with the following conditions: 94°C for 20 s, 95°C for 3 s, 60°C for 30 s, for 40 cycles, then 95°C for 15 s and 60°C for 1 min. Assays were performed in triplicate, with two biological replicates. Primer sequences are included in Supplementary Table 1. Data were normalised to *ACT1* and analysed with QuantStudio Design & Analysis Software.

### Killing assay

Overnight cultures of *C. albicans* were subcultured for 3 h prior to the assay. Yeast cells were resuspended in RPMI-1640 containing 2% pooled human plasma, 1× PSQ and 0.05 μg/mL of DOX, then incubated with primary neutrophils at an MOI of 5 at 37 °C with 5% CO₂. Fungal killing was assessed using two methods.

For CFU determination, after 2 h incubation, cell suspensions were collected, vortexed to disrupt clumps, serially diluted, and plated on YPD agar. Plates were incubated at 30 °C for 48 h to determine viable colonies. Percentage killing was calculated as follows: [1 - (CFU of neutrophils treated group/CFU of *C. albicans* only control)] x 100.

For metabolic viability, after 2.5 h incubation, samples were treated with 0.1% Trition X-100 to lyse neutrophils, washed with by 1X PBS twice and incubated with 1X Alamar Blue for 16 h at 37°C. FLUOstar Omega Microplate Reader (BMG LABTECH) was used to measure fluorescence.

### Growth kinetics

*C. albicans* overnight cultures were diluted in YPD media supplemented with 0.05 μg/mL of DOX to a starting OD_600_ of 0.02 in an Eppendorf tube. Cultures (100 μL) were transferred into a clear flat-bottom 96-well plate, then sealed with Breathe-Easy® sealing membrane (Diversified Biotech BEM-1). The OD_600_ was read by a microplate reader (BMG Labtech) every 15 min for 20 h at 30°C and 700 RPM shaking.

### Filamentation assay

Overnight cultures were subcultured and grown in YPD for 3 h. Prior to being diluted to an OD_600_ of 0.2 in RPMI-1640 supplemented with 0.05 μg/mL of DOX and 10% human pooled plasma and incubated for 2.5 h at 37°C. An Olympus IX70 microscope was used to take images of each strain.

### ROS production assay

Overnight cultures of *C. albicans* strains were subcultured for 3 h in YPD medium supplemented with 0.05 μg/mL DOX at 30 °C, while shaking at 200 RPM. Following opsonisation, yeast cells were resuspended in PBS containing 10% pooled human plasma and 0.05 μg/mL DOX, and ROS production was assayed using a luminol assays as previously described^50^. In brief, the yeast suspension was added at MOI=5 to white, flat-bottom 96-well plates (Thermo Fisher Scientific) containing 100 μL neutrophils resuspended in ROS medium [HBSS (with Ca²⁺ and Mg²⁺), 1× HEPES, and 0.025% (v/v) human serum albumin (HSA; Sigma-Aldrich)]. Fluorescence was recorded every 2.5 min over a 4 h period using a BMG FLUOstar plate reader (excitation 490 nm, emission 515 nm).

### Phagocytosis assay

Human neutrophils were seeded into glass-bottom dishes in NETs medium (RPMI1640 supplemented with 0.05 μg/mL DOX, 0.025% HSA and 10mM HEPES) and allowed to adhere for 15 min at 37 °C. Mid-log phase *C. albicans* (OD_600_ 0.7–0.8) were washed, resuspended in NETs medium, and added at MOI 3 for 30 min to allow phagocytosis. Extracellular fungi were stained with 10 μg/mL CFW (Sigma-Aldrich) on ice, followed by fixation with 4% paraformaldehyde and staining with Alexa Fluor 488–phalloidin (Thermo Fisher). Samples were imaged using a Leica LASX workstation (40X oil objective). At least 50 neutrophils per replicate were analysed. Phagocytosed fungal cells were defined as CFW-negative cells, and the phagocytic index was calculated as the number of intracellular fungi per neutrophil^51,52^. Images were processed using Fiji (ImageJ).

### NET formation assay

Human neutrophils were co-cultured with *C. albicans* (MOI 5, pre-opsonised in 10% human plasma) in NETs medium on poly-L-lysine–coated coverslips in 24-well plates. After 4 h incubation at 37 °C with 5% CO₂, samples were fixed with 4% paraformaldehyde, permeabilised with 0.1% Triton X-100, and blocked in 1% BSA with 0.05% Triton X-100. NETs were stained using a mouse anti-DNA/histone complex antibody (3D9 antibody, Max Planck Institute, Berlin^35^), followed by Alexa Fluor 488-conjugated donkey anti-mouse IgG (Thermo Fisher). Coverslips were mounted with ProLong Gold Antifade containing DAPI ( Invitrogen) and imaged after 24 h. Images were acquired using a Leica SP8 confocal microscope and analysed with Fiji (ImageJ). At least seven fields of view were analysed per replicate. Unstimulated neutrophils were included as controls.

### ELISAs

Sub-cultured *C. albicans* strains were prepared in advance, diluted to 2×10^7^ cells/mL to achieve an MOI of 5. Isolated neutrophils were resuspended in RPMI-1640 supplemented with 10% FCS and Penicillin/Streptomycin/Glutamine (PSQ, Thermo Fisher Scientific), and stimulated with *C. albicans* and 100ng/mL bacterial lipopolysaccharide (LPS from *E. coli* O127:B8, Sigma Aldrich). Samples were incubated overnight at 37°C with CO_2_. Supernatant was collected and frozen at - 20°C for future use. IL-8 concentration was measured using a DuoSet ELISA kit as per manufacturer’s instructions (R&D Systems). Absorbance was measured at 450 nm and 540 nm using a BMG FLUOstar plate reader, and a standard curve was generated to obtain the cytokine concentration.

### Neutrophil RNA extraction and RT-qPCR

Subcultured *C. albicans* strains were prepared and diluted to achieve an MOI of 5. Isolated neutrophils were resuspended in RPMI-1640 supplemented with 10% FCS and PSQ and stimulated with 100 µL yeast suspension for 90 min at 37 °C with 5% CO₂. Following incubation, plates were placed on ice, the supernatant was removed, and cells were lysed in 350 µL RLT buffer (Qiagen). RNA was extracted according to the manufacturer’s instructions and quantified using a NanoDrop spectrophotometer. cDNA was synthesised using a first-strand synthesis kit with random primers (Promega) according to the manufacturer’s protocol. Quantitative PCR was performed using SYBR Green master mix (Thermo Fisher Scientific), with gene-specific primers on a QuantStudio system. Primer sequences are included in Supplementary Table 2. Cycling conditions were 95 °C for 20 s, followed by 40 cycles of 95 °C for 3 s and 60 °C for 30 s, with a melt curve step.

### Immunofluorescent staining of cell wall components

Overnight *C. albicans* cells were subcultured to mid-log phase from a starting OD_600_ of 0.1 and grown for 3 h at 30°C, 200 RPM. The cell suspension was then collected and fixed with 4% PFA on ice for 30 min. Fixed cells were washed with PBS and stained in 96-well plates using the following probes: (1) β-1,3-glucan was labelled with 5 µg/mL Dectin-1-Fc (R&D Systems), followed by goat anti-human IgG conjugated to Alexa Fluor 488 (Thermo Fisher Scientific), (2) chitin was stained with 10 µg/mL Calcofluor White (Sigma), (3) mannan was detected with 50 µg/mL TRITC-conjugated concanavalin A (Thermo Fisher Scientific). All incubations were performed on ice or at room temperature in the dark for 1 h, with washes in FACS buffer (1× PBS, 1% FCS, 0.5 mM EDTA) between steps. Stained cells were analysed by Novocyte 3000 ACEA flow cytometer (Biosciences), with 30, 000 events observed. The MFI was quantified.

Following immunofluorescent staining for exposed β-1,3-glucan, total mannan and chitin, yeast cells were imaged using a Leica LASX live cell imaging workstation with Photometrics Prime 95B sCMOS camera with 100X lens. Images acquisition was performed using a Leica SP5 confocal microscope with 63X lens.

### Quantification of cell wall polysaccharides by high-pressure-ion-chromatography (HPIC)

Cell wall polysaccharides composition was determined by HPIC measurement of the hydrolysis products of the purified cell walls form yeast cell populations^53^. For GRACE strain samples yeast cells were grown overnight in YPD in the presence of DOX at 30°C. Washed yeast cells were disrupted in a FastPrep machine (MP Biomedicals) and the lysate centrifuged and the pellet washed with 1 M NaCl. Samples were boiled for 10 min at 100 °C in SDS extraction buffer (500 mM Tris-HCl pH7.5, 2% (w/v) SDS, 0.3 M β-mercaptoethanol, 1 mM EDTA), and then freeze dried. The glucan, mannan and chitin levels were determined by measurement of glucose, mannose and glucosamine, after hydrolysis in 2 M trifluoroacetic acid at 100 °C for 3 h. Hydrolysates were analysed as before^54^. Samples were measured in a Dionex carbohydrate analyser using a CarboPac PA20 column (0.4×150mm), guard column and an ED50 Pulsed amperometric detector (PAD). Samples were eluted using 5-100 mM gradient, flow rate 0.008 ml/min for 25 min.

## Statistical analysis

Data were analysed using GraphPad Prism software (v10). All results are presented as mean ± standard error of the mean (SEM) from a minimum of three independent biological replicates (n≥3). Comparisons between two groups were performed using Student’s t-test, while multiple group comparisons were analysed using one-way ANOVA with Dunnett’s post-hoc test for comparisons against the control. Statistical significance was defined as P<0.05.

## Author contributions

**HJ** performed experiments and analysed data; **IL** performed HPIC experiments and analysed data; **AN**, **NG** and **SD** supervised fungal experiments; **BA** conceived and supervised the study. **HJ** and **BA** wrote the manuscript.

All authors read, provided input and approved the final manuscript.

## Data availability

Raw data is available upon request from the lead and corresponding authors.

## Acknowledgments

We thank all the blood donors for participating in our study. We acknowledge University of Bristol’s Flow Cytometry Facility and Wolfson Bioimaging Facility. For the purpose of open access, the author has applied a ‘Creative Commons Attribution (CC BY) licence to any Author Accepted Manuscript version arising from this submission.’

---

**Fig. S1.**
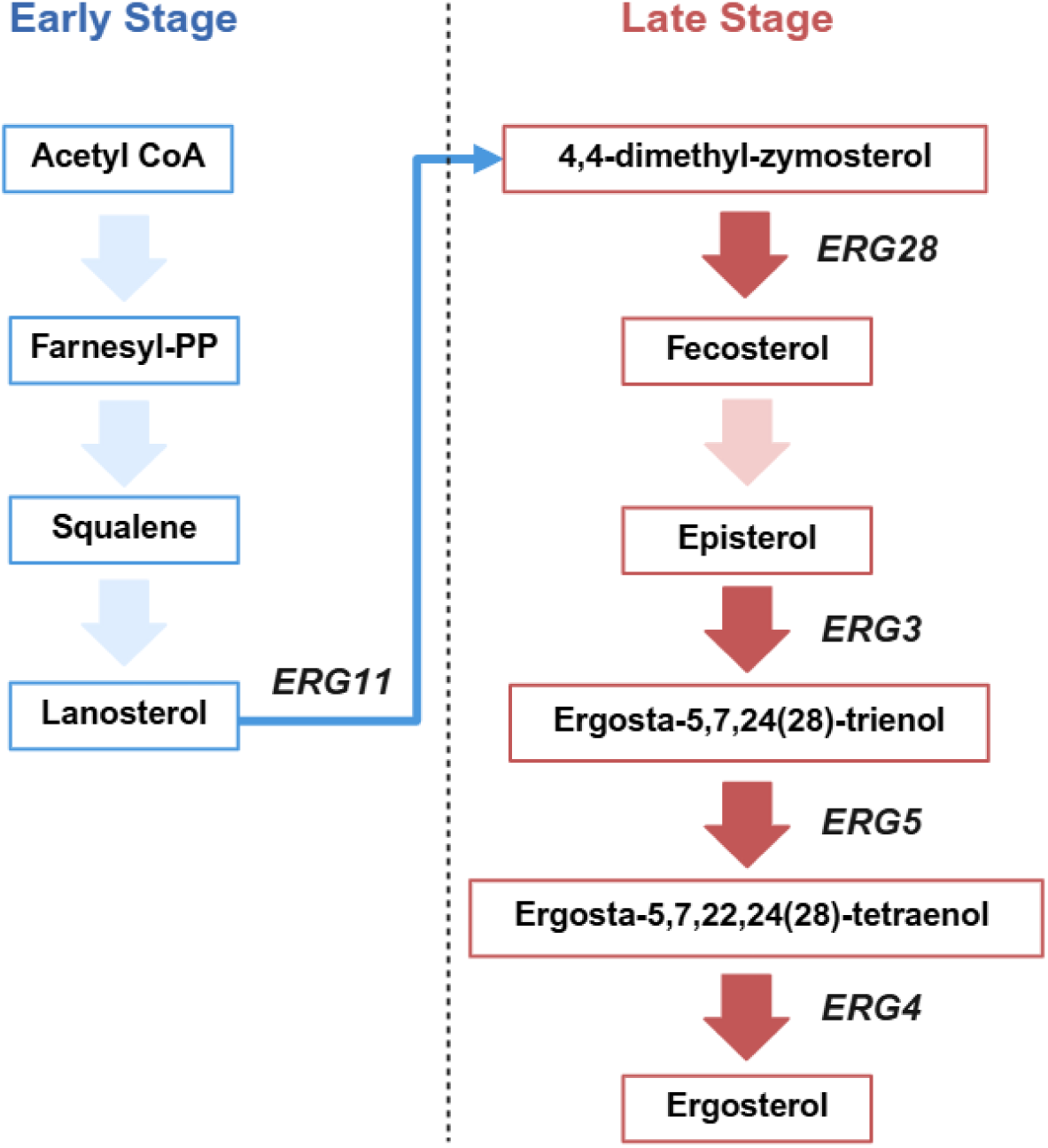
Schematic overview of the ergosterol biosynthesis pathway in *C. albicans*. The pathway can be divided into early (blue) and late (red) stages^28,32,38^. Genes investigated in this study are indicated in bold: *ERG28*, *ERG3*, *ERG5*, and *ERG4*. Created with BioRender.com.

**Fig. S2.**
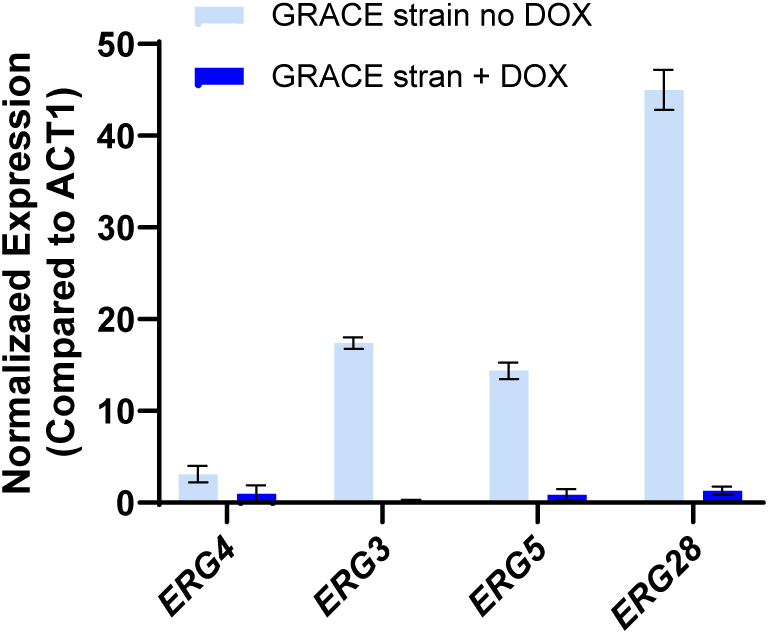
Quantitative RT-PCR analysis of gene expression. *C. albicans* strains were incubated overnight for 16 h in YPD medium supplemented with 0.05 μg/mL DOX, followed by RNA extraction. Target gene transcript levels were calculated using the 2^−ΔΔCT^ method and then normalised to the housekeeping gene *ACT1*. Representative figure from one of two biological replicates.

**Fig S3.**
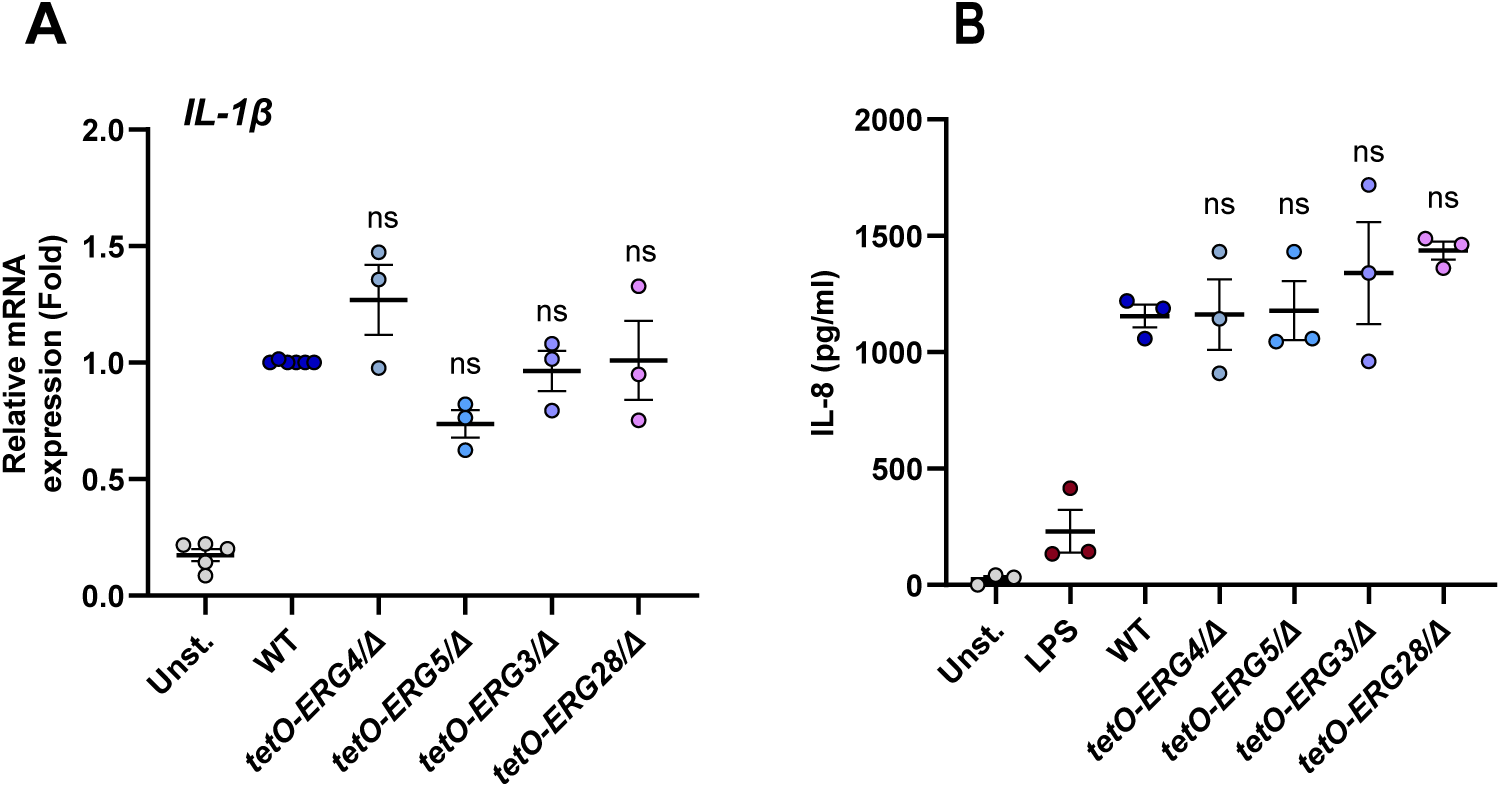
The *ERGs* conditional mutant strains do not affect neutrophil cytokine and chemokine production. **(A)** Human peripheral blood neutrophils were stimulated with WT, *ERG4-*, *ERG5-*, *ERG3-*, and *ERG28*-depleted *C. albicans* cells at an MOI of 5 for 90 min. The pellet was collected and *IL-1*β mRNA abundance was assessed by qRT-PCR after normalising to GAPDH (n=3-6 blood donors). **(B)** Neutrophils were cultured with WT and mutant strains at an MOI of 5 for 5 h, the supernatant was collected and IL-8 protein abundance was assessed by ELISA (n=3 blood donors). Error bars represents mean ± SEM. Significance was calculated using one-way ANOVA with Dunnett’s multiple comparison post-hoc test. ns: *P* > 0.05.

## Supplemental tables

**Supplementary table 1:**
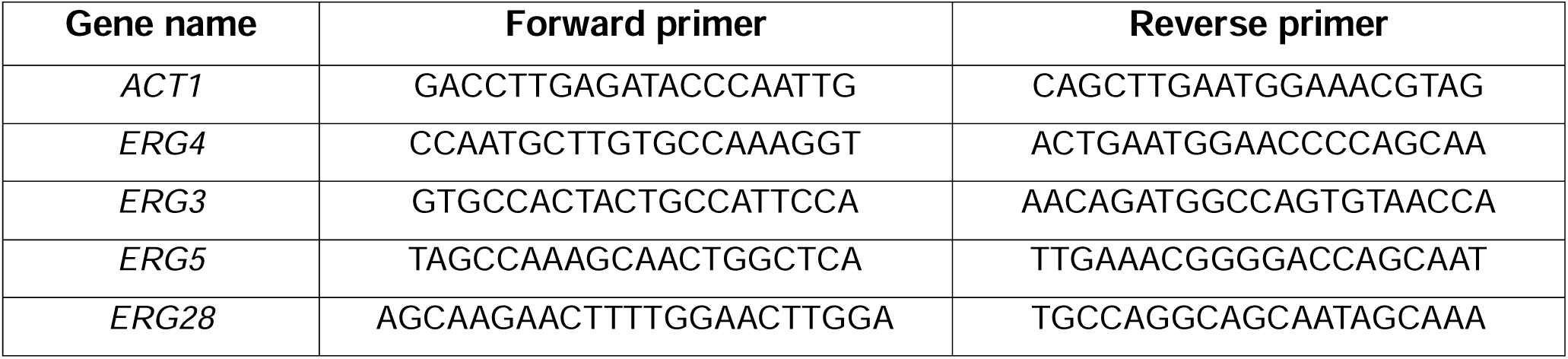
Primers for *C. albicans*.

**Supplementary table 2:**
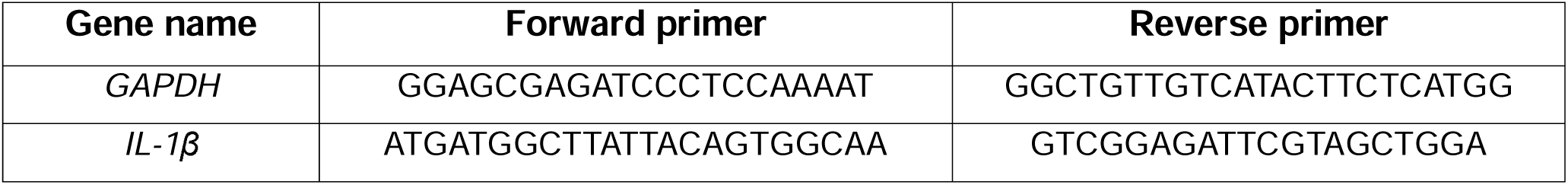
Primers for human neutrophils.

